# A quorum sensing-regulated type VI secretion system containing multiple nonredundant VgrG proteins is required for interbacterial competition in *Chromobacterium violaceum*

**DOI:** 10.1101/2022.04.29.490089

**Authors:** Júlia A. Alves, Fernanda C. Leal, Maristela Previato-Mello, José F. da Silva Neto

## Abstract

The environmental pathogenic bacterium *Chromobacterium violaceum* kills Gram-positive bacteria delivering violacein packed into outer membrane vesicles, but nothing is known about its contact-dependent competition mechanisms. In this work, we demonstrate that *C. violaceum* utilizes a type VI secretion system (T6SS) containing multiple VgrG proteins primarily for interbacterial competition. The single T6SS of *C. violaceum* contains six *vgrG* genes, which are located in the main T6SS cluster and four *vgrG* islands. Using T6SS-core component null mutant strains, western blot, fluorescence microscopy, and competition assays, we show that the *C. violaceum* T6SS is active and required for competition against Gram-negative bacteria such as *Pseudomonas aeruginosa* but dispensable for *C. violaceum* infection in mice. Characterization of single and multiple *vgrG* mutants revealed that, despite having high sequence similarity, the six VgrGs show little functional redundancy, with VgrG3 showing a major role in T6SS function. Our coimmunoprecipitation data support a model of VgrG3 assembling heterotrimers with the other VgrGs. Moreover, we determined that the promoter activities of T6SS genes increased at high cell density, but the produced Hcp protein was not secreted under such condition. This T6SS growth-phase-dependent regulation was dependent on CviR but not on CviI, the components of a *C. violaceum* quorum sensing (QS) system. Indeed, a Δ*cviR* but not a Δ*cviI* mutant was completely defective in Hcp secretion, T6SS activity, and interbacterial competition. Overall, our data reveal that *C. violaceum* relies on a QS-regulated T6SS to outcompete other bacteria and expand our knowledge about the redundancy of multiple VgrGs.

**IMPORTANCE:** The type VI secretion system (T6SS) is a contractile nanomachine used by many Gram-negative bacteria to inject toxic effectors into adjacent cells. The delivered effectors are bound to the components of a puncturing apparatus containing the protein VgrG. The T6SS has been implicated in pathogenesis and, more commonly, in competition among bacteria. *Chromobacterium violaceum* is an environmental bacterium that causes deadly infections in humans. In this work, we characterized the single T6SS of *C. violaceum* ATCC 12472, including its six VgrG proteins, regarding its function and regulation. This previously undescribed *C. violaceum* T6SS is active, regulated by QS, and required for interbacterial competition instead of acute infection in mice. Among the VgrGs, VgrG3, encoded outside of the main T6SS cluster, showed a major contribution to T6SS function. These results shed light on a key contact-dependent killing mechanism used by *C. violaceum* to antagonize other bacteria.

## INTRODUCTION

Bacteria engage in complex ecological relationships with other organisms and very often compete with other bacteria. Interbacterial competition can occur at a distance through diffusible secreted inhibitory factors or mediated by contact (1-4). Contact-mediated competition involves the injection of toxins directly into a target cell through complex protein machineries called secretion systems, such as the type IV, type V, and type VI secretion systems (3, 5). The type VI secretion system (T6SS), found in many Gram-negative bacteria, delivers toxic effectors into both bacterial and host eukaryotic cells (6-8). However, its main function is to mediate interbacterial competition, affecting the dynamics of bacterial communities in multiple ecological contexts, such as during interactions between bacterial pathogens and the microbiota (4, 9). In addition to hijacking host cellular processes and intoxicating competing bacteria, novel roles have been described for the T6SS and its effectors, including antifungal activity, metal acquisition, resistance against amoeba predation, and several intraspecies social interactions (9-12).

The T6SS is a complex nanomachine resembling the contractile tail of bacteriophages. It is composed of at least 14 core components that are assembled in subcomplexes: a cytoplasmic baseplate, an intermembrane complex, a contractile sheath-like structure (VipAB), and an inner tube of polymerized Hcp sharpened in its tip by VgrG trimers and PAAR proteins (13-16). The translocation of toxic protein effectors by the T6SS directly inside a target cell involves the firing of a puncturing apparatus (Hcp/VgrG/PAAR) by contraction of the VipAB sheath (17). The ATPase ClpV recycles the sheath components, which are then reused for new T6SS firing events (13, 18). The effectors can be “specialized” when they are a domain covalently fused to the expelled components VgrG, PAAR or Hcp, or “cargo” effectors, which comprise independent proteins interacting directly or via adaptor proteins with VgrG, PAAR or Hcp (15, 19, 20). VgrG (valine-glycine repeat protein G) proteins have a modular architecture containing a core gp27– gp5 portion with puncturing and structural function and sometimes an additional C-terminal region serving as an effector or adaptor domain (6, 15, 21, 22). Even in bacteria harboring a single T6SS, the presence of multiple VgrG proteins encoded by genes located in the main T6SS cluster and *vgrG* islands is common (23). However, little is known about the redundancy of multiple VgrGs for the function of the T6SS (23-26).

The expression and activity of T6SSs can be tightly regulated at several levels in response to different environmental cues, such as reactive oxygen species, metal scarcity, temperature, pH, and membrane damage caused by the attack of competitor bacteria. For bacteria having more than one T6SS, each of them can be differentially regulated to exert a particular function (12, 16, 27, 28). Among the multitude of regulatory mechanisms controlling T6SS expression, regulation by quorum sensing (QS) systems is a recurrent theme with very distinct outcomes (27). For instance, the QS regulation of T6SSs has been described in several bacteria, such as *Pseudomonas aeruginosa* (29), *Vibrio cholerae* (30), *Burkholderia thailandensis* (31), *Aeromonas hydrophila* (32), and *Vibrio fluvialis* (33).

*Chromobacterium violaceum* is a free-living Gram-negative bacterium found in the soil and water of tropical and subtropical regions. Despite its free-living lifestyle, *C. violaceum* causes systemic infections in humans and animals by employing several virulence factors, such as a Cpi-1/1a T3SS, which injects effectors into host eukaryotic cells (34-37). *C. violaceum* produces violacein, a violet hydrophobic compound with broad-spectrum biocide activity that targets the cytoplasmic membranes of Gram-positive bacteria (38, 39). By delivering violacein packed into outer membrane vesicles (OMVs), *C. violaceum* outcompetes Gram-positive but not Gram-negative bacteria in a contact-independent manner (40, 41). In addition to violacein, bacteria of the genus *Chromobacterium* antagonize other organisms, such as pathogenic fungi, parasites, and larvae of disease-transmitting insects, by using extracellular secreted molecules, such as chitinases, proteases, hydrogen cyanide, and romidepsin (42-45). Most of these compounds are specifically produced at high cell density via activation by the QS system CviI/CviR, a circuit composed of the regulator CviR, and by CviI, an enzyme that produces homoserine lactone (HSL)-type autoinducers (46-49). However, the role of contact-dependent mechanisms for *C. violaceum* competition remains largely unknown.

In this study, we used interbacterial competition assays to reveal that *C. violaceum* employs its single T6SS to overcome competitor Gram-negative bacteria, such as *P. aeruginosa*. Virulence assays with a murine infection model indicated that *C. violaceum* T6SS mutants were as virulent as the wild-type strain. These data indicate that the *C. violaceum* T6SS is more relevant to interbacterial competition than to bacterial pathogenesis. Furthermore, we demonstrate that the *C. violaceum* T6SS is active, expressed under laboratory conditions, and regulated by cell density and QS. The study of the six *vgrG* genes found in the *C. violaceum* genome revealed little functional redundancy, given that VgrG3 showed a preponderant role in relation to the other five VgrGs in the assembly of a functional T6SS with antibacterial activity. Coimmunoprecipitation assays indicated that VgrG3 interacts directly with other VgrGs *in vivo*, suggesting that VgrG3 could nucleate the formation of VgrG heterotrimers to allow the secretion of effectors bound to the other VgrGs.

## RESULTS

### *C. violaceum* has a single T6SS containing multiple VgrG proteins

We searched for T6SS genes in the genome of *C. violaceum* ATCC 12472 (50) using BLAST searches with conserved T6SS core components (VipA and Hcp) and the TXSScan tool (51). These *in silico* analyses retrieved a single T6SS whose genes are located in a main cluster containing 31 open reading frames (from CV_3963 to CV_3991) organized as two divergently transcribed large putative operons (Fig. 1A). This T6SS main cluster contains all genes encoding the T6SS core components (*tssA*-*tssM* and *PAAR*) found in a single copy, except for *vgrG* (*tssI*), which was found in two copies (*vgrG*1 and *vgrG2*). There is also a *tagF* gene, described as a posttranslational inhibitor of T6SS activity in other bacteria (16). Downstream of both *vgrG*1 and *vgrG2*, there are genes encoding putative effectors (the phospholipases of the Tle1 family with DUF2235 domain, CV_3990 and CV_3971) (52), chaperone/adaptor proteins (Tap-1, DUF4123) (53, 54), and putative immunity proteins (DUF3304) (Fig. 1A). Using *vgrG1* and *vgrG2* for BLAST searches, we detected four additional *vgrG* genes (*vgrG3, vgrG4, vgrG5*, and *vgrG6*) dispersed in other regions of the *C. violaceum* genome. Each such *vgrG* is located in small orphan VgrG clusters that contain genes encoding putative effectors, such as Rhs-repeat-containing proteins (CV_1429, CV_1431, CV_1238, and CV_1239), Tle5 (CV_1234) and Tle1 (CV_0012) phospholipases, and a protein with lysozyme-like and peptidase M23 domains (CV_0024) (Fig. 1A). These analyses indicate that *C. violaceum* has a single T6SS and a repertoire of potential T6SS-delivered effectors encoded by genes located near those encoding six VgrG proteins.

**Fig 1.**
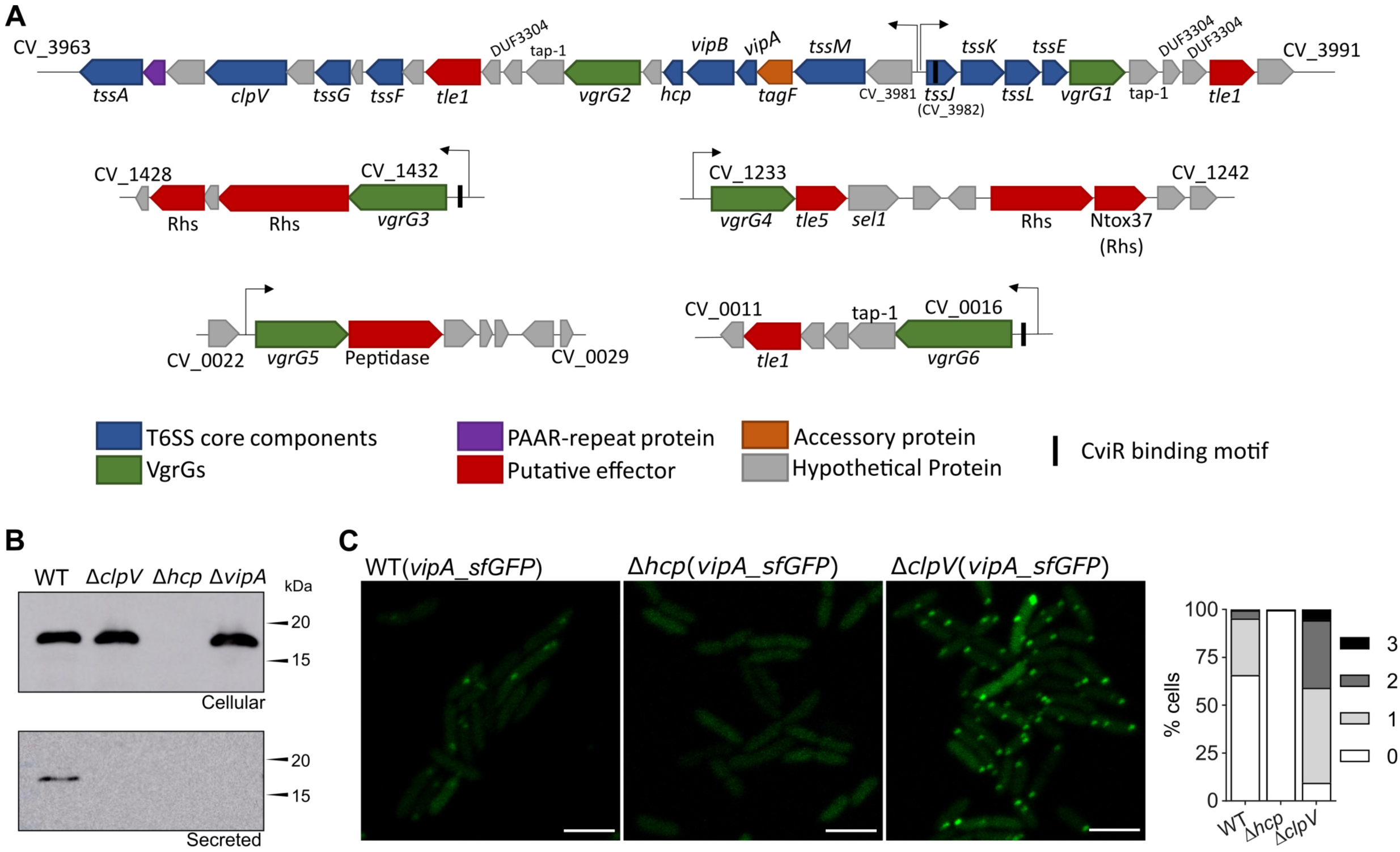
The single T6SS of *C. violaceum* is active and contains multiple VgrG proteins. (A) The genes encoding the T6SS components are located in a main T6SS cluster and four minor orphan VgrG clusters in the genome of *C. violaceum* ATCC 12472. Conserved T6SS components and effectors are colored according to their functions. The six VgrGs are indicated in green. The arrows above the genes indicate promoter regions. Genes of the main T6SS are organized into two putative large operons transcribed from divergent promoters (regions between CV_3981 and *tssJ*). (B) Secretion of Hcp depends on T6SS core components. Western blot of Hcp with an anti-Hcp antibody from the *C. violaceum* WT and the indicated T6SS null mutant strains using whole cellular extracts or cell-free supernatants. (C) Activity of the T6SS visualized by fluorescence microscopy of VipA_sfGFP. Assembled and contracted sheaths appear as GFP foci. The foci were counted manually, and the percentages indicate the proportions of bacterial cells with one, two or three GFP foci at the same time. Scale = 3 µm.

### The *C. violaceum* T6SS is functional and active under laboratory growth conditions

To verify the functionality of the *C. violaceum* T6SS, we analyzed the expression and secretion of Hcp by immunoblotting assays with an anti-Hcp serum developed against the purified His-Hcp protein. In the cellular fraction, Hcp was detected from LB-grown log-phase cultures of WT, Δ*clpV*, and Δ*vipA* strains (Fig. 1B). However, Hcp was detected in the cell-free supernatants of the WT but not the Δ*clpV* and Δ*vipA* strains, indicating that the T6SS core components VipA (sheath structure) and ClpV (sheath depolymerization) are required for Hcp secretion. As expected, Hcp was not detected in either the cellular or secreted fractions of the Δ*hcp* mutant strain (Fig. 1B). To assess the dynamics of the T6SS sheath *in vivo*, we constructed a plasmid-borne carboxy-terminal fusion of the VipA protein with the superfolded green fluorescent protein (sfGFP) (17), and the resulting construct was transferred to the WT and T6SS mutants. Visualization by fluorescence microscopy in the WT strain showed a high activity of the T6SS sheath in log-phase cultures, as shown by the high number of sfGFP foci (Fig. 1C). As expected, no sfGFP focus was observed in Δ*hcp* (Fig. 1C) since Hcp polymerization is necessary for sheath assembly. Furthermore, the images suggest that nondynamic foci accumulated in Δ*clpV*, possibly because, in the absence of ClpV, the VipAB sheath structure remained assembled after the firing event (Fig. 1C). These results indicate that the *C. violaceum* T6SS is functionally active and requires the T6SS core components Hcp, VipA, and ClpV for its proper function.

### The T6SS contributes to interbacterial competition but not to virulence in *C. violaceum*

Considering that T6SSs mediate bacterial competition and are involved in bacterial pathogenesis (4, 8, 55, 56), we tested whether the T6SS is involved in such processes in *C. violaceum* (Fig. 2). To verify the competitive capacity of *C. violaceum* against a broad range of bacteria, we performed a visual screening that consisted of the coculture of *C. violaceum* WT or T6SS mutants (purple due to violacein) with phylogenetically distinct bacteria (Fig. 2A). *C. violaceum* WT was able to overcome almost all targeting bacteria (purple output), except for *Enterobacter cloacae* ATCC 13047 (Ecl) and *P. aeruginosa* PAO1 (Pao) (Fig. 2A). However, the *C. violaceum* advantage was lost for the Δ*clpV* and Δ*hcp* mutant strains in cocultures against *Escherichia coli* (Ec), *P. aeruginosa* ATCC 27853 (Pa), and *Stenotrophomonas maltophilia* (Sma) and to a lesser extent against *Salmonella* Typhimurium (St), *Klebsiella pneumoniae* ATCC 13883 (Kp), and *Shigella sonnei* (Ss) (Fig. 2A). The T6SS mutant strains grew similarly to the WT strain in LB medium (Fig. S1A), suggesting that their decreased competitive fitness is due to the loss of the T6SS functionality itself instead of a general fitness impairment.

**Fig 2.**
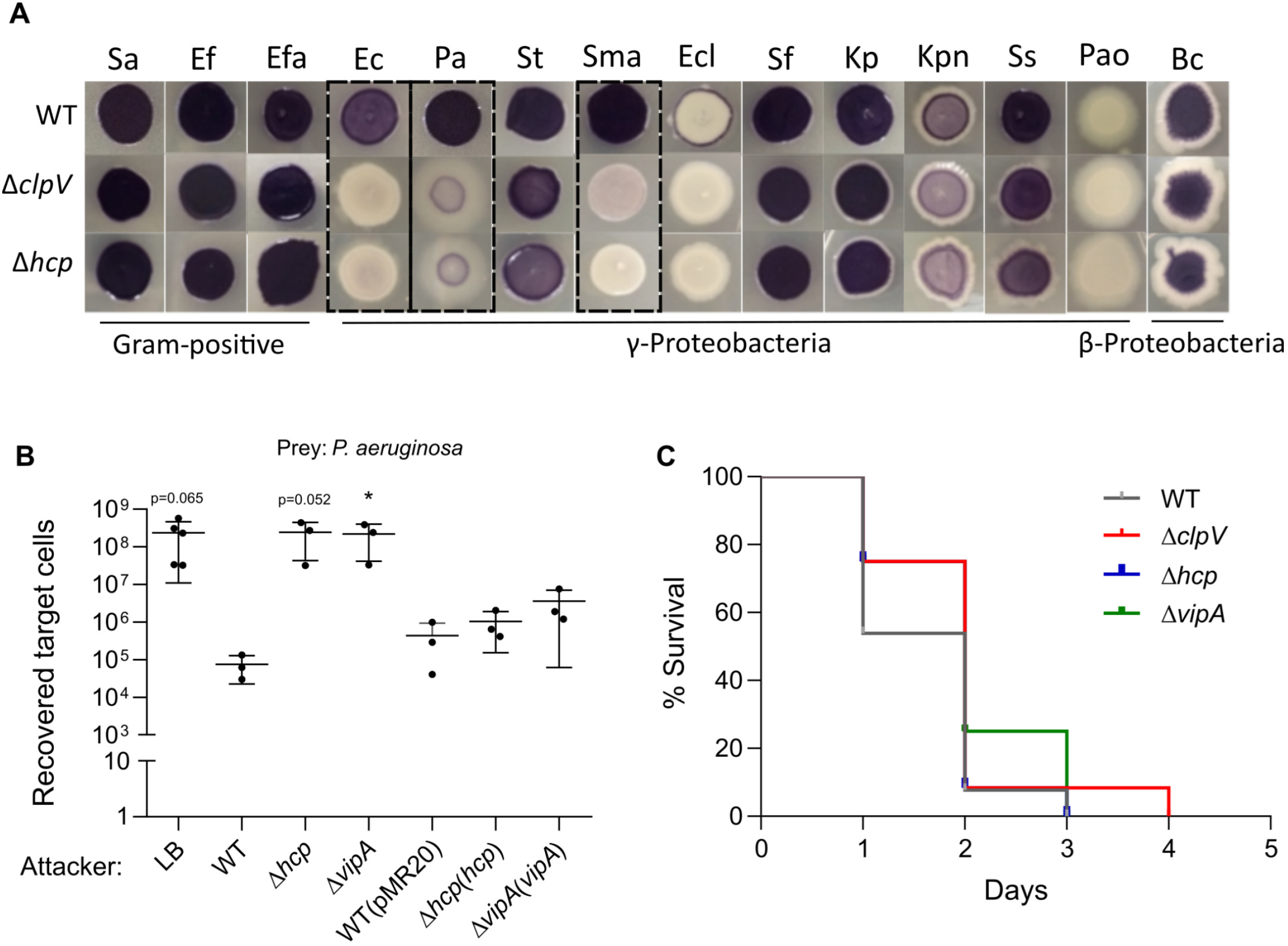
The T6SS contributes to interbacterial competition but not to virulence in *C. violaceum*. (A) *C. violaceum* outcompetes many Gram-negative bacteria via the T6SS. Qualitative growth competition assay of *C. violaceum* WT or T6SS mutants against the indicated bacteria (details in Table S3). Predominantly purple spots (due to violacein production by *C. violaceum*) indicate the cocultures in which *C. violaceum* was able to outcompete the target bacteria. The *C. violaceum* T6SS showed a major role in competition against *E. coli* (Ec), *P. aeruginosa* ATCC 27853 (Pa) and *S. maltophilia* (Sma). (B) *C. violaceum* employs its T6SS to kill *P. aeruginosa*. A quantitative antibacterial assay was performed by counting the CFU of *P. aeruginosa* ATCC 27853 without (LB) or after coculture with *C. violaceum* WT, T6SS mutants (Δ*hcp* and Δ*vipA*) or complemented strains. Data are from at least three independent biological assays. Bars show the mean and SD. Statistical analyses were performed using the unpaired Student’s t test, and asterisks indicate variations in relation to the WT. *p < 0.05. (C) The T6SS does not contribute to *C. violaceum* virulence. BALB/c mice were intraperitoneally infected with 10^6^ CFU of the indicated *C. violaceum* strains. The animals were monitored for five days (n=13 for WT; n=12 for Δ*clpV*; n=11 for Δ*hcp*; and n=4 for Δ*vipA*). Statistical analysis by the log-rank (Mantel-Cox) test showed no significant difference among the strains.

To further confirm the role of the T6SS as an antibacterial weapon, we performed a quantitative antibacterial assay using cocultures of *C. violaceum* (attacker) with *P. aeruginosa* ATCC 27853 (prey) at a ratio of 5:1 (Fig. 2B). The recovery of *P. aeruginosa* was at least three logs higher in competition against the T6SS mutants than against the WT strain, indicating *C. violaceum* T6SS-mediated killing activity. In fact, the prey recovery competing against the T6SS mutants was at the same levels as the negative control using LB medium instead of *C. violaceum*. The complemented strains killed *P. aeruginosa* similarly to the WT strain (Fig. 2B). Together, these results indicate that the T6SS is a killing machinery that provides a competitive advantage for *C. violaceum* against other bacteria, likely helping it persist in mixed bacterial populations. The role of the T6SS in virulence was assessed using a mouse model of acute infection (57). After intraperitoneal injection, animals inoculated with three distinct T6SS mutants succumbed as fast as those inoculated with the WT strain (Fig. 2C), indicating that the T6SS does not contribute to *C. violaceum* virulence, at least under our tested conditions.

### Among the six VgrGs, VgrG3 has major roles in T6SS assembly and function

The protein VgrG is the sole T6SS core component whose genes were found in multiple copies in the *C. violaceum* genome (Fig. 1A). The six VgrG proteins from *C. violaceum* showed high sequence identity (70 to 93%) and a similar domain organization (Fig. S2), raising the question of whether they would have redundant roles. To address this question, we constructed null mutant strains with individual or sequential deletion of the *vgrG* genes and a sextuple mutant (Δ*vgrG1-6*) with each single *vgrG* gene reintroduced using a plasmid. The *vgrG* mutant strains showed no growth impairment, as assessed by growth curves (Fig. S1B and C). These strains were used to investigate the roles of the six VgrGs in the activity/assembly (Fig. 3) and antibacterial function (Fig. 4) of the T6SS.

**Fig 3.**
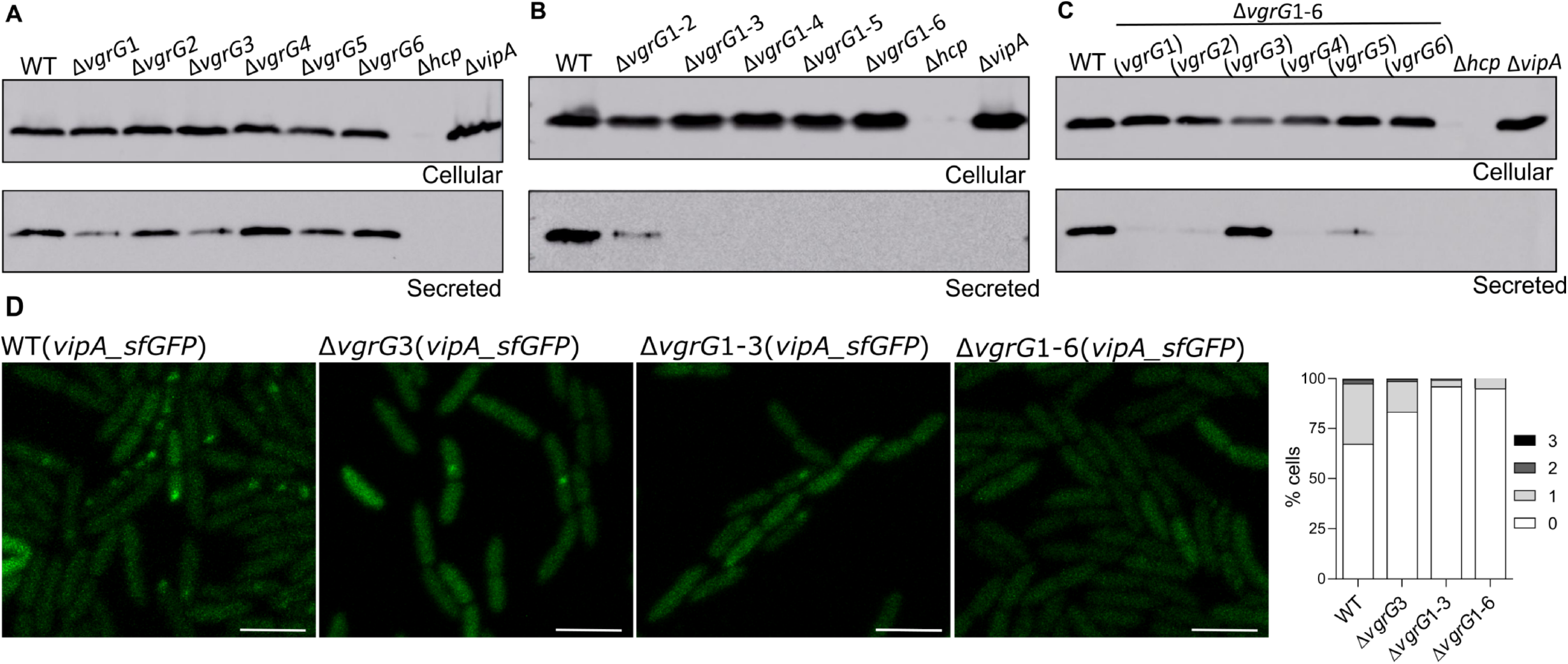
Each VgrG distinctly affects T6SS activity, but VgrG3 has a preponderant role. (A-C) Analysis of VgrG-dependent activity of the T6SS evaluating Hcp secretion. Western blot with anti-Hcp to evaluate Hcp expression (cellular) and secretion (secreted) in (A) individual *vgrG* mutants, (B) sequential *vgrG* mutants or (C) the sextuple Δ*vgrG1-6* mutant harboring each single *vgrG* gene. (D) Effects of VgrGs on the assembly and activity of the T6SS visualized by the formation of VipA_sfGFP foci. The GFP foci were observed by fluorescence microscopy and counted in the indicated *vgrG* mutant strains. Scale = 3 µm.

**Fig 4.**
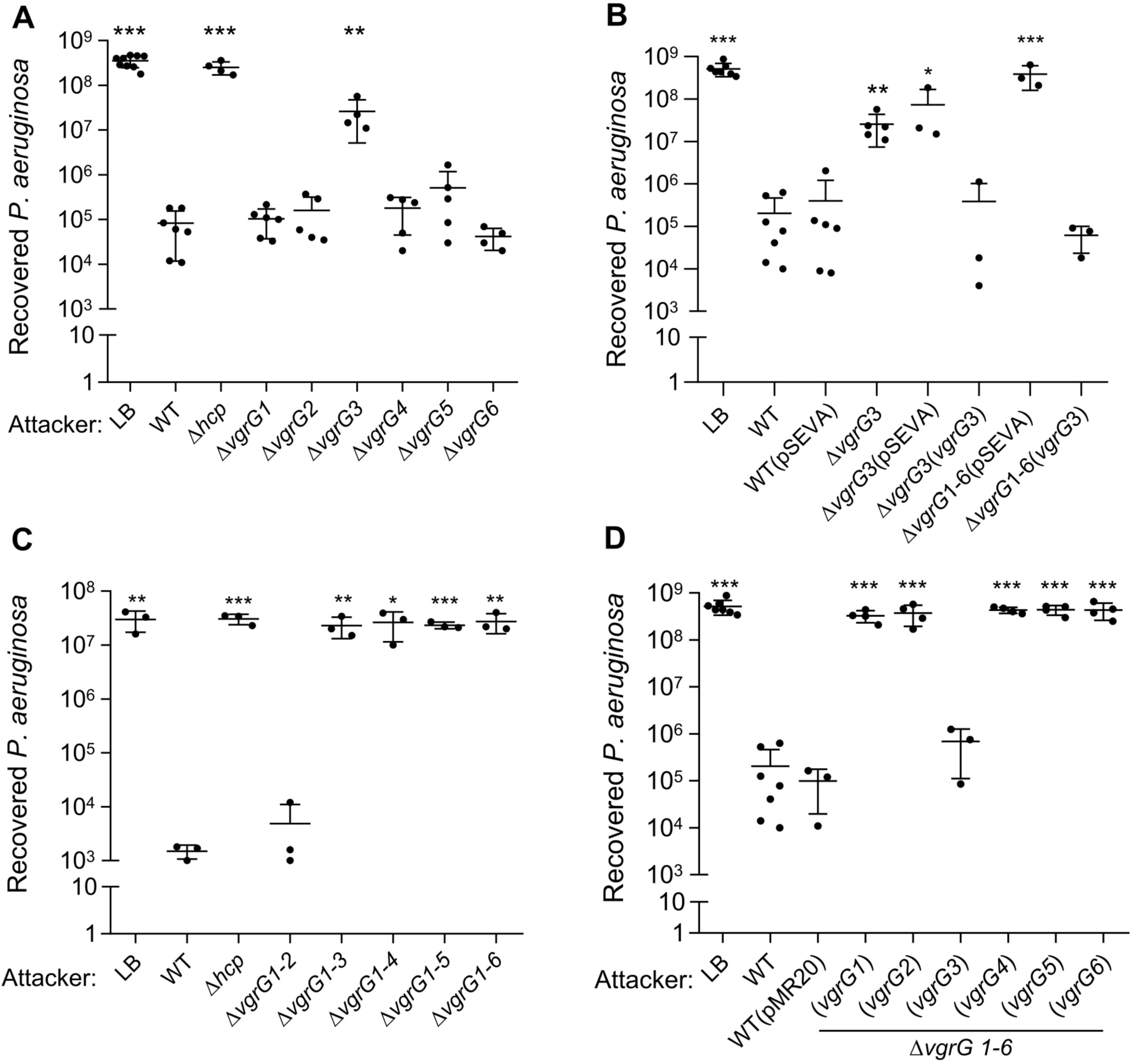
VgrG3 is the most important VgrG for *C. violaceum* antibacterial activity. A quantitative antibacterial assay was performed by counting the CFU of *P. aeruginosa* ATCC 27853 without (LB) or after coculture with *C. violaceum*. Competitions were performed using (A) the individual *vgrG* mutants, (B) the complemented Δ*vgrG*3(*vgrG3*) strain, (C) the sequential *vgrG* mutants, and (D) the sextuple Δ*vgrG1-6* mutant harboring each single *vgrG* gene. Each point represents biological replicates of distinct assays. Bars show the mean and SD. Statistical analyses were performed using the unpaired Student’s t test, and asterisks indicate variations in relation to the WT. *p < 0.05, **p < 0.02, ***p < 0.001.

The immunoblotting of secreted Hcp from log-phase cultures revealed that the individual deletion of each *vgrG* had a distinct impact on the T6SS functionality, with marked decreases in Hcp secretion in the Δ*vgrG1* and Δ*vgrG3* mutants and slight or no effects in the other *vgrG* mutants (Fig. 3A). The double mutant Δ*vgrG1-2* showed residual secretion of Hcp, but no Hcp secretion was detected from the triple to the sextuple *vgrG* mutants, indicating that the combined deletion of *vgrG1, vgrG2*, and *vgrG3* resulted in the complete loss of T6SS functionality (Fig. 3B). Immunoblotting assays using the sextuple mutant with the reintroduction of each single *vgrG* gene indicated that the presence of *vgrG3* fully restored the ability of the Δ*vgrG1-6* mutant to secrete Hcp, while the presence of the other five individual *vgrG* genes resulted in very weak Hcp secretion (Fig. 3C). As expected, the Hcp protein was no longer detected in the Δ*hcp* mutant (cellular or supernatant fractions) or in the supernatant of the Δ*vipA* mutant (Fig. 3A, B, and C). These results indicate that VgrG3 has a preponderant role over the other VgrGs in Hcp secretion and suggest that VgrG3 is able to form functional homotrimers in the absence of the other VgrGs. The role of VgrG3 in T6SS functionality was further confirmed by fluorescence microscopy using VipA_sfGFP. The number of sfGFP foci decreased in the Δ*vgrG*3 mutant compared to those found in the WT strain and was almost abolished in the Δ*vgrG1-3* and Δ*vgrG1-6* mutant strains (Fig. 3D).

To verify the contribution of each VgrG to interbacterial competition, we performed competition assays against *P. aeruginosa* ATCC 27853 using our panel of *C. violaceum vgrG* mutants (Fig. 4). Among the single *vgrG* mutants, only *ΔvgrG3* showed a decreased competitive fitness compared to that of the WT strain, although at levels lower than the control (LB) and the Δ*hcp* mutant (Fig. 4A). This phenotype was fully reverted in the complemented strain *ΔvgrG3*(*vgrG3*) and was unaffected in the strains with the empty vector (Fig. 4B). Regarding the sequential *vgrG* mutants, Δ*vgrG1-2* displayed no change in competitive capacity, but Δ*vgrG1-3* and all other multiple *vgrG* mutants showed strong impairments in competition, similar to that found for the Δ*hcp* mutant (Fig. 4C). The complete loss of competitiveness of the sextuple mutant Δ*vgrG1-6* was fully reverted when this strain was provided with the *vgrG*3 gene, but no effect was observed using the other *vgrG* genes (Fig. 4D). Altogether, these data demonstrate that VgrG3 is the most important VgrG for the antibacterial activity of the *C. violaceum* T6SS, probably owing to its important structural role for the overall T6SS function.

### VgrG3 interacts directly with other VgrGs *in vivo*

Given the main role of VgrG3 in T6SS assembly and function, we hypothesize that VgrG3 could interact with other VgrGs to allow the delivery of their cognate effectors. To test this hypothesis, we developed a construct into the pMR20 vector (*vgrG*3-HA) to express VgrG3 fused to a hemagglutinin epitope at its C-terminus (VgrG3-HA) for coimmunoprecipitation assays (Co-IP) using a monoclonal anti-HA antibody (Fig. 5). After introduction of this construct into the Δ*vgrG*3 mutant, we confirmed by immunoblotting that the VgrG3-HA protein was expressed (Fig. 5A) and able to complement the competitive deficiency of this mutant strain against *P. aeruginosa* ATCC 27853 (Fig. 5B). Co-IP assays were performed from the Δ*vgrG3*(*vgrG3*-HA) strain and the WT(pMR20) strain (a negative control), and the immunoprecipitated proteins were identified by mass spectrometry (Table S4). Excluding many abundant cellular proteins detected in the Co-IPs from both strains, such as ribosomal proteins (Table S4), six proteins were detected specifically from the Δ*vgrG3*(*vgrG3*-HA) strain (Hcp, VgrG1, VgrG2, VgrG3, VgrG5, and CV_2846) (Fig. 5C). Therefore, the Co-IP data showed that VgrG3 interacts with VgrG1, VgrG2 and VgrG5 *in vivo*, most likely forming heterotrimers.

**Fig 5.**
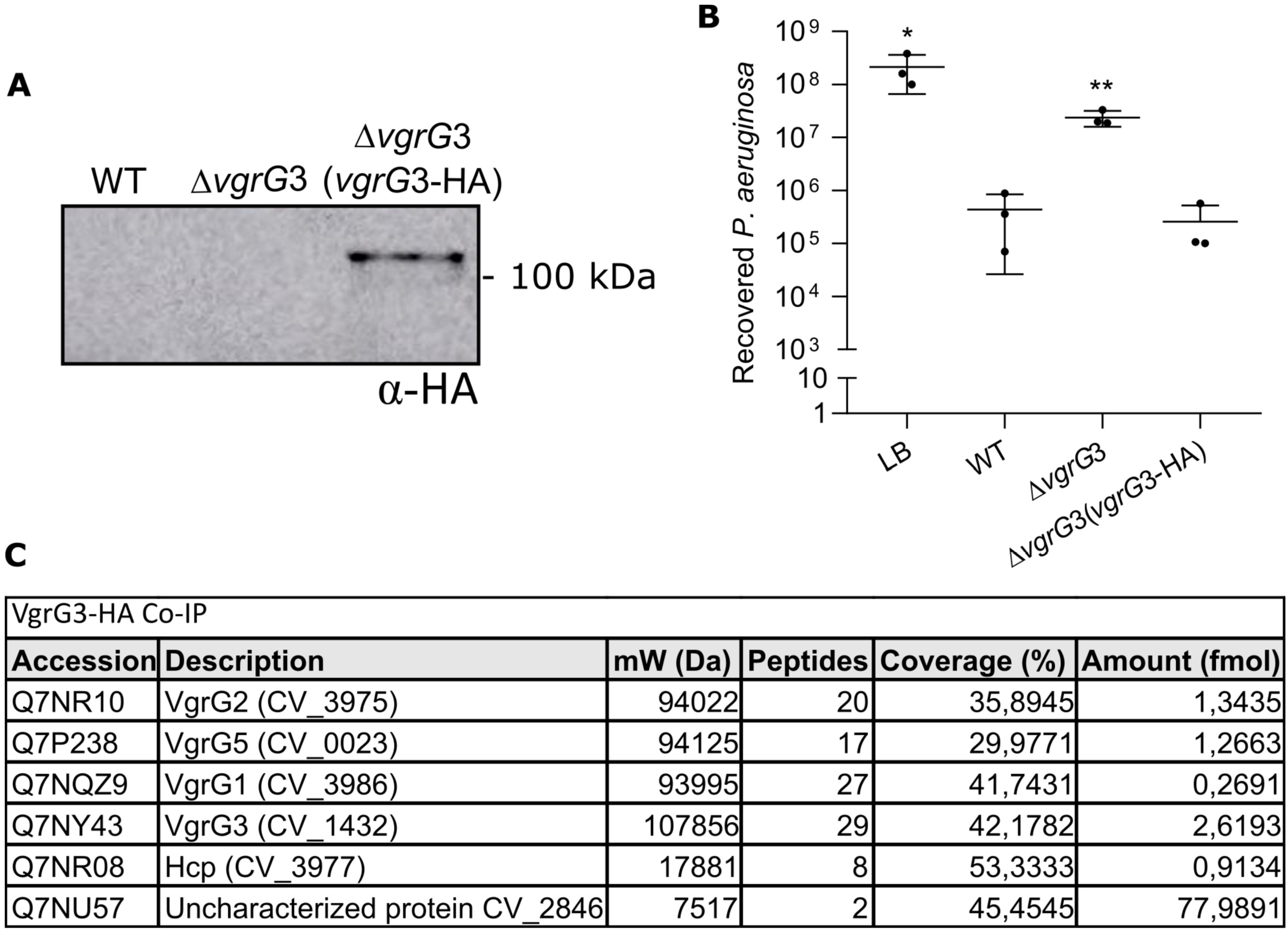
VgrG3 interacts directly with other VgrGs *in vivo*. (A) Confirmation of VgrG3-HA expression. Western blot with a monoclonal anti-HA antibody revealed VgrG3-HA expression (predicted molecular weight of 110 kDa) in the Δ*vgrG*3 strain harboring a vector expressing the construct *vgrG*3-HA. (B) VgrG3-HA functionally complements the Δ*vgrG*3 mutant strain. Interbacterial competition of the *C. violaceum* indicated strains against *P. aeruginosa* ATCC 27853. Competitive fitness was rescued in the Δ*vgrG*3(*vgrG*3-HA) strain. Bars indicate the mean and SD. Statistical analyses were performed using the unpaired Student’s t test, and asterisks indicate variations in relation to the WT. *p < 0.05, **p < 0.02. (C) Several VgrGs coimmunoprecipitated with VgrG3-HA. Proteins identified after mass spectrometry of coimmunoprecipitation with VgrG3-HA. As a control, Co-IP was also performed with the WT(pMR20) strain. Many abundant cellular proteins detected in the Co-IP from both strains (Table S4) may represent contaminants.

### Regulation of the *C. violaceum* T6SS is growth phase-dependent and affected by CviR

It has been demonstrated in other bacteria that the T6SS is regulated by QS (27, 30, 32, 33). The HSL-based QS circuit CviI/CviR controls many processes in *C. violaceum* (46-49). Considering the presence of three predicted CviR binding sites associated with T6SS genes in *C. violaceum* (47) (Fig. 1A), we hypothesize that T6SS expression is under QS regulation. To test this hypothesis, we measured the activity of T6SS promoters along growth curves by β-galactosidase activity assays using transcriptional reporter fusions (Fig. 6). The expression levels of the two divergent promoters that drive transcription of the two operons in the main T6SS cluster (Fig. 1A) increased throughout the growth curve in the WT strain but to a lesser extent in Δ*cviR*, indicating that CviR activates these promoters mainly at high cell density (Fig. 6A and B). With respect to the six *vgrG* genes, the regions upstream of *vgrG1, vgrG2*, and *vgrG6* showed no promoter activity regardless of the growth phase, while the promoters of *vgrG*3, *vgrG*4 and, at lower levels, *vgrG*5 were active and showed a slight tendency to be more expressed at high cell density than at low cell density (Fig. 6C). The absence of activity for the *vgrG1* and *vgrG2* fusions and the intraoperonic locations of *vgrG1* and *vgrG2* suggest that these genes lack their own promoters and are transcribed from the promoters of the two divergent operons. Detailed analysis of the two more active *vgrG* promoters, *vgrG3* and *vgrG4*, revealed that their expression levels were markedly decreased in the Δ*cviR* mutant strain at all points along the growth curve (Fig. 6D and E), indicating a full dependence on CviR. Overall, these data show that the QS regulator CviR exerts transcriptional activation on many T6SS genes in *C. violaceum*.

**Fig 6.**
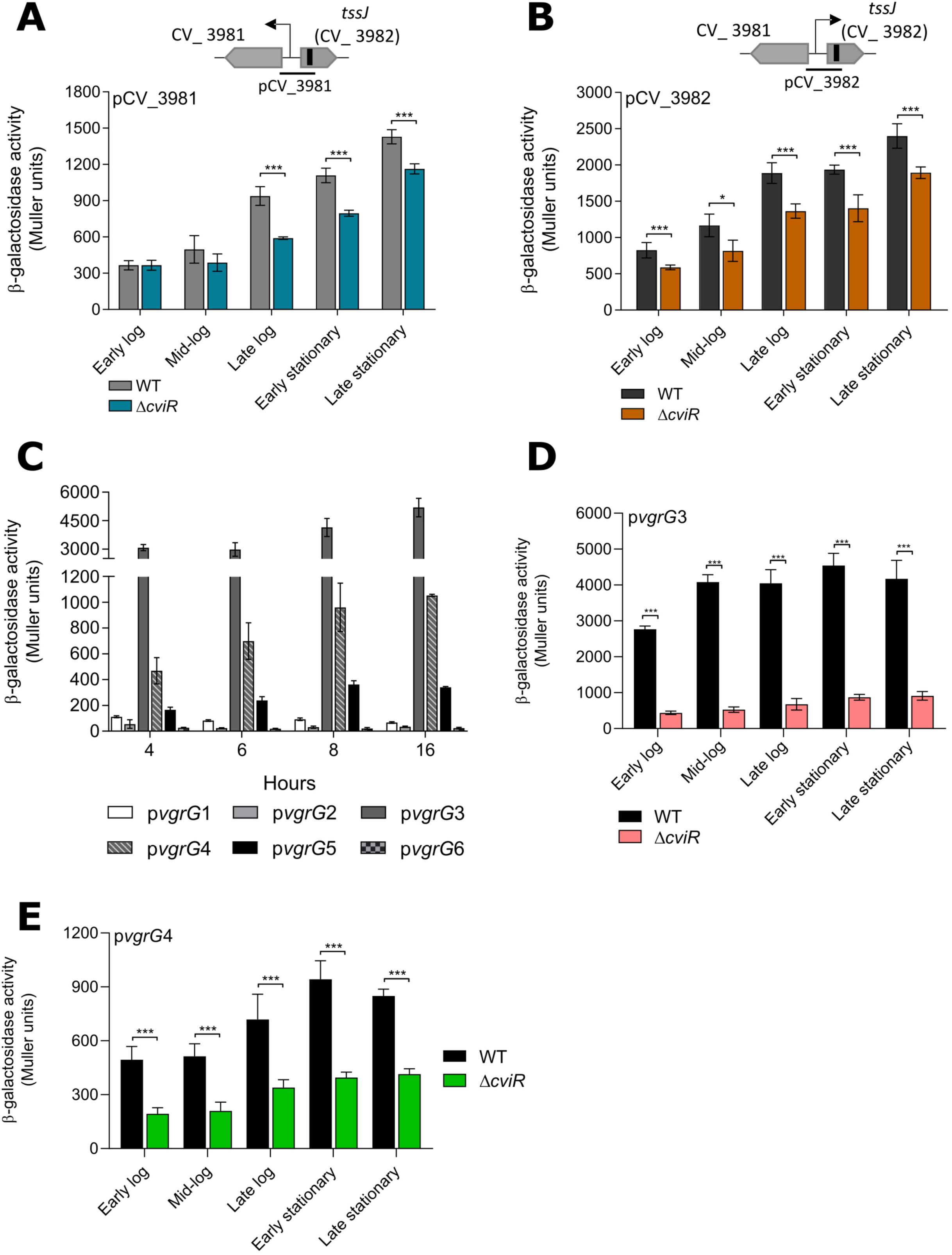
The promoter activity of T6SS genes increases at high cell density and is activated by the QS regulator CviR in *C. violaceum*. (A-E) β-Galactosidase activity assays of the indicated promoters were performed along the indicated time points of the growth curve. The strains (WT or Δ*cviR*) harboring the *lacZ* fusions were cultivated in liquid LB from an OD_600_ of 0.01 until the indicated growth phases. (A and B) Promoters pCV_3981 and pCV_3982, driving the expression of the two divergent operons of the main T6SS cluster. (C) Promoters of the six *vgrG* genes along the growth curve. (D and E) Effects of CviR on the promoter activity levels of *vgrG3* and *vgrG4*. Bars show the mean and SD. *p < 0.05, **p < 0.02, ***p < 0.001 (Unpaired Students’s t test).

Considering the CviR-dependent transcriptional regulation of T6SS genes, we tested the effects of CviR and CviI on T6SS functionality (Fig. 7). To determine whether T6SS activity was dependent on cell density, Hcp expression and secretion throughout the growth curve were verified by immunoblotting (Fig. 7A). In the WT strain, although the Hcp protein was highly expressed throughout the growth curve (cellular fraction), its secretion occurred at low but not at high cell density (Fig. 7A), suggesting that *C. violaceum* inhibits T6SS activity at high cell density by a posttranslational mechanism. Surprisingly, Hcp expression and secretion were markedly impaired in the Δ*cviR* strain but barely affected in the Δ*cviI* strain, independent of the growth phase (Fig. 7A). Consistently, in fluorescence microscopy assays with VipA_sfGFP, the GFP foci were abolished in the Δ*cviR* strain but almost unaffected in the Δ*cviI* strain (Fig. 7B). Similarly, in competition assays against *P. aeruginosa* ATCC 27853, the Δ*cviR* mutant showed a complete loss of competitive capacity, as observed for the Δ*hcp* mutant, while the Δ*cviI* mutant killed *P. aeruginosa* as efficiently as the WT strain (Fig. 7C). Altogether, these results indicate that *C. violaceum* depends on CviR but not on CviI for the assembly of a functional T6SS, suggesting that the QS system CviI/CviR regulates the T6SS by a noncanonical mechanism.

**Fig 7.**
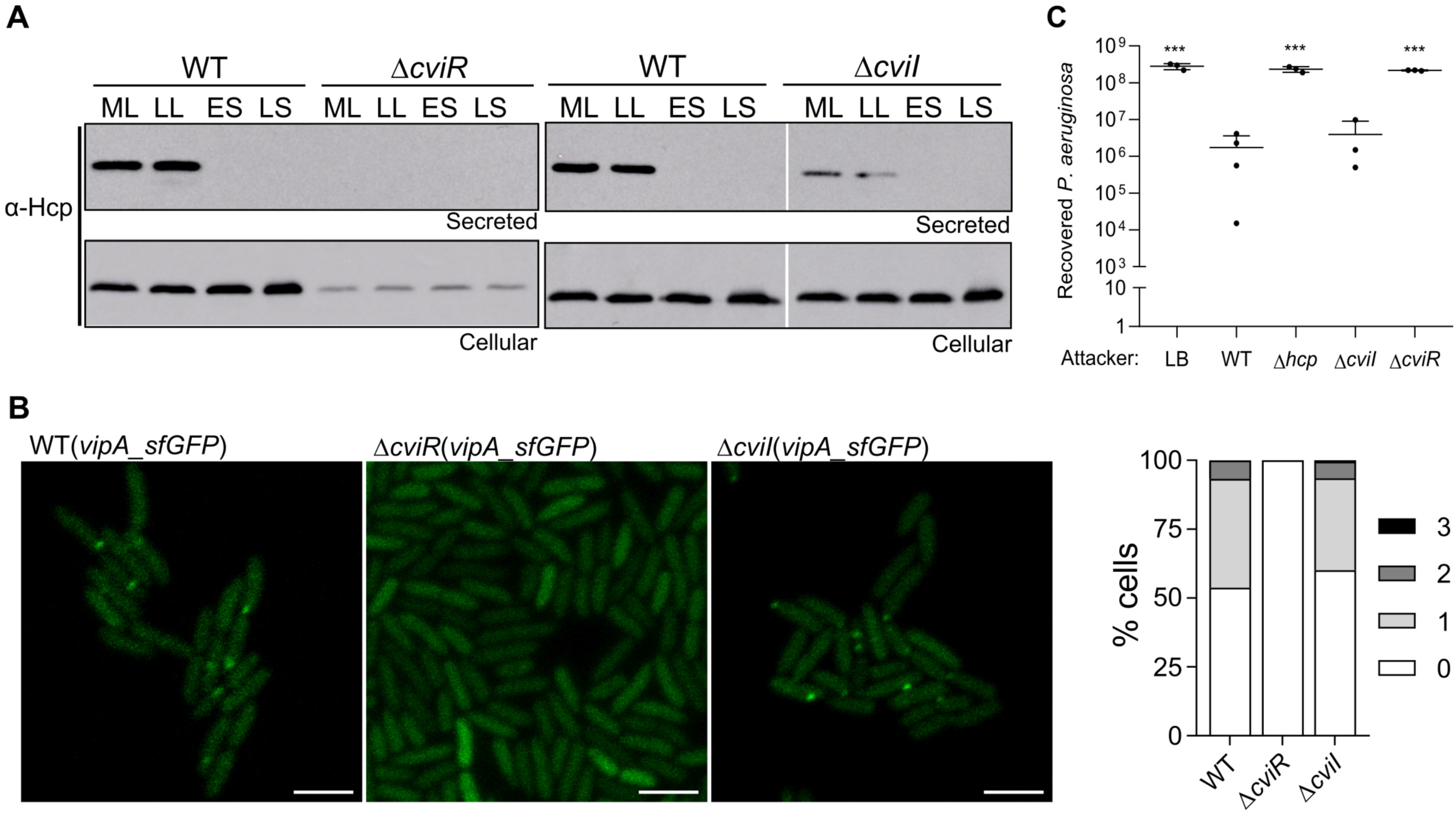
*C. violaceum* depends on CviR but not on CviI for the assembly of a functional T6SS. (A) Effects of cell density, CviR and CviI on Hcp production and secretion. Western blot of Hcp in cellular and secreted fractions of WT, Δ*cviR*, and Δ*cviI* strains at different points of the growth curve: ML, mid-log phase; LL, late log phase; ES, early stationary; LL, late stationary. (B) Fluorescence microscopy showing T6SS sheath assembly through the formation of VipA_sfGFP foci. Scale = 3 µm. The graph shows the average number of foci per cell. (C) Recovery of *P. aeruginosa* ATCC 27853 after 4 hours in coculture with the indicated *C. violaceum* strains. Bars indicate the mean and SD. Statistical analyses were performed using the unpaired Student’s t test, and asterisks indicate variations in relation to the WT. ***p < 0.001.

## DISCUSSION

The T6SS has been characterized in detail in some model bacteria. Studying this system in several different bacterial species allows us to gain novel insights regarding its function and regulation and find new families of antibacterial and anti-eukaryotic T6SS effectors (8, 16, 20). In this work, we characterized the function and regulation of the T6SS of *C. violaceum*, an environmental pathogenic bacterium. Our results show that the single T6SS of *C. violaceum* is active, regulated by QS and has a major role in contact-dependent interbacterial competition. These findings add the T6SS to the already known armamentarium used by *C. violaceum* to antagonize other bacteria, which includes diffusible molecules and violacein delivered into OMVs (36, 38, 40, 41, 49). Given that the T6SS mutants showed no virulence attenuation and the predicted T6SS effectors seem to be antibacterial toxins, it is tempting to speculate that *C. violaceum* relies on its Cpi-1/1a T3SS but not on its T6SS to inject effectors that affect key functions in host mammalian cells (34, 37).

The presence of six VgrG proteins in *C. violaceum* showing high sequence identity and no toxic domains gave us the opportunity to study the functional redundancy of VgrGs for T6SS function. These proteins are encoded by *vgrG* genes located in the main T6SS cluster and four *vgrG* islands and surrounded by putative T6SS effector genes (encoding phospholipases, Rhs-repeat-containing proteins, and a putative peptidase), a genomic organization commonly found in other bacteria (23, 26, 52, 58). Moreover, all six VgrGs presented a similar domain organization: a core gp27-gp5 portion followed by an additional DUF2345 C-terminal domain, which has been found in many VgrGs (25) and contributes to the VgrG/effector interaction (21). VgrG3 and VgrG4 also presented an additional C-terminal unstructured region after the DUF2345 domain. However, we provide experimental evidence, using single and multiple *vgrG* mutant strains, that in *C. violaceum* the VgrGs showed little functional redundancy, with the VgrG3 protein playing a more important role in relation to the other five VgrGs to the T6SS function: (i) among the single *vgrG* mutants, *ΔvgrG3* showed a marked decrease in Hcp secretion and loss of competitive fitness against *P. aeruginosa*; (ii) only *vgrG3* among the six *vgrG* genes fully restored the ability of the Δ*vgrG1-6* mutant to secrete Hcp and to kill *P. aeruginosa*; (iii) the *vgrG3* promoter was the most active among the *vgrG* promoters and showed full dependence on the QS regulator CviR; and (iv) VgrG3 interacted *in vivo* with other VgrGs (VgrG1, VgrG2, and VgrG5), most likely favoring the assembly of VgrG heterotrimers to allow the secretion of effectors associated with each VgrG, as described for the three VgrGs of *V. cholerae* (59). Distinct results, indicative of greater functional redundancy, have been described in other bacteria harboring multiple VgrGs. For instance, in *Burkholderia cenocepacia* (ten VgrGs), *Serratia marcescens* (two VgrGs), and *Agrobacterium tumefaciens* (four VgrGs), the individual deletion of any *vgrG* had no impact on Hcp secretion (23, 25, 58). Conversely, in *A. hydrophila* and *V. cholerae* (each with three VgrGs), the individual deletion of specific *vgrGs* abolished Hcp secretion (59, 60). As expected, single-copy T6SS core components such as Hcp, VipA, and ClpV were absolutely required for the T6SS functionality in *C. violaceum*.

Our data indicate that the *C. violaceum* T6SS is an offensive system because it is active even in laboratory conditions (as indicated by the high number of sfGFP foci and the easy detection of Hcp in supernatants of log-phase liquid cultures) and efficiently killed many Gram-negative bacteria regardless of whether they had a T6SS to a counterattack. This offensive pattern, described in other bacteria such as *S. marcescens* (61), differs from the defensive strategy employed by *P. aeruginosa*, where the T6SS assembly and firing occurs only in response to an incoming attack and involves complex posttranslational regulation by the threonine phosphorylation pathway (TPP) (16, 62). Although the *C. violaceum* T6SS can be considered an offensive system, our results indicate that Hcp secretion ceased at high cell density despite the normal production of Hcp and the maximum expression of the T6SS promoters under such condition via activation by the QS regulator CviR. Similarly, a T6SS2 of *V. fluvialis* was regulated by QS (33), and Hcp secretion was repressed at high cell density (63). *C. violaceum* seems to lack the TPP pathway but harbors TagF encoded in the main T6SS cluster; TagF is a protein that can block T6SS activity independent of TPP in other bacteria (16, 61, 64). We speculate that *C. violaceum* posttranslationally blocks T6SS firing events in nonpolymicrobial cultures at high cell density by activating *tagF* together with the other T6SS genes from a CviR-dependent promoter. This strategy ensures that the T6SS components will be produced but that the T6SS firing will be blocked, avoiding physical damage and activation of stress responses in nearby kin cells inhabiting densely populated cultures. These adventitious events have been described in recipient cells targeted by T6SSs at least during interspecies competition (65, 66). Many processes regulated by QS in *C. violaceum* depend on both CviI and CviR, the sole QS circuit described in this bacterium (46, 47, 49). Our results indicate a clear role of CviR but not CviI in the T6SS function, suggesting that CviR could act independently of the HSL autoinducers produced by CviI. Future works investigating the global impact of CviI and CviR on the *C. violaceum* transcriptome will contribute to revealing whether this QS system operates together with other QS circuits to control the T6SS.

## MATERIALS AND METHODS

### Bacterial strains, plasmids, and growth conditions

The bacterial strains and plasmids used in this work are listed in Table S1. *E. coli* and *C. violaceum* strains were grown with agitation at 37°C in Luria-Bertani (LB) medium or on LB agar plates (LB supplemented with 15 g/L agar). When necessary, media were supplemented with antibiotics at the following concentrations: gentamycin 40 μg/mL, tetracycline 5-10 μg/mL, and kanamycin 50 μg/ml for *C. violaceum*; and gentamycin 20 μg/mL, tetracycline 12 µg/mL, kanamycin 50 μg/ml, and ampicillin 100 μg/mL for *E. coli*.

### Construction of the *C. violaceum* mutant and complemented strains

Strains harboring in-frame deletions of the selected T6SS genes were constructed by an allelic exchange protocol, as described previously (40, 67, 68). Briefly, the flanking regions of the selected genes were amplified by PCR with specific primers (Table S2) and cloned into the suicide vector pNPTS138. After electroporation into *E. coli* S17-1, the constructs were transferred by conjugation to *C. violaceum* ATCC 12472. Transconjugants were selected on LB agar with kanamycin 50 μg/mL and ampicillin 100 μg/mL. Isolated colonies were grown in liquid LB followed by counterselection on LB plates with 16% sucrose. For the multiple *vgrG* mutants, the procedure was repeated sequentially, starting from the Δ*vgrG1* mutant to the final sextuple mutant Δ*vgrG1-6*. The null mutant strains were confirmed by PCR using specific primers (Table S2). To complement the T6SS mutants, the respective T6SS genes were amplified by PCR with specific primers (Table S2) and cloned into the replicative plasmids pMR20 or pSEVA. The resulting constructs were transferred to the respective *C. violaceum* mutant strain by conjugation. Similarly, each *vgrG* gene cloned into pMR20 was also introduced into the sextuple mutant Δ*vgrG1-6*.

### Interbacterial competition

For an initial screening, attacker *C. violaceum* strains were competed against several prey bacteria (Table S3) and analyzed for a purple output by visual inspection (indicating *C. violaceum* overgrowth). Attacker and prey bacteria were grown overnight on LB agar plates, resuspended in LB at an optical density at 600 nm (OD_600_) of 1 and mixed at a ratio of 1:1. The mixtures (5 µL) were spotted on LB agar plates and incubated at 37°C overnight prior to image acquisition. For the quantitative competition assay of *C. violaceum* strains against *P. aeruginosa* ATCC 27853, bacterial strains grown overnight on LB agar were resuspended in liquid LB and normalized to an OD_600_ of 5. *C. violaceum* WT and T6SS mutant strains (attacker) or LB (control) were mixed with the target bacterium at a proportion of 5:1. The mixtures (10 µL) were spotted on LB agar to allow contact-dependent competition. After 4 hours at 37°C, the bacterial spots were recovered and resuspended in 1 mL of LB. Serial dilutions were plated on LB agar with kanamycin to allow *P. aeruginosa* CFU counting.

### Mouse virulence assay

*C. violaceum* virulence in mice was assayed using an intraperitoneal (i.p.) infection model, as previously described (57). Briefly, six-week-old BALB/c female mice were challenged with i.p. injection of 10^6^ CFU of *C. violaceum* WT and mutant strains. To check the injected dose, serially diluted bacterial samples were plated on LB agar for CFU counting. The survival of infected mice was monitored for five days. All experiments using mice were performed in the Animal Facility of the Faculdade de Medicina de Ribeirão Preto (FMRP-USP) according to the ARRIVE guidelines, and protocol number 037/2018 was approved by the local animal ethics committee (CEUA, FMRP-USP). The CEUA follows the Ethical Principles in Animal Research adopted by the National Council for the Control of Animal Experimentation (CONCEA).

### β-galactosidase activity assay

The promoters of the two large operons located in the main T6SS cluster (divergent regions between CV_3981 and CV_3982, *tssJ*) and the promoter regions of each *vgrG* were amplified by PCR (Table S2), cloned into the pGEM-T easy plasmid (Promega), and subcloned into the pRK*lacZ*290 vector to generate transcriptional fusions to the promoterless *lacZ* gene. The resulting constructs were introduced into *C. violaceum* wild-type and Δ*cviR* strains. To assess the expression of these promoters according to the cell density, the strains were diluted to an OD_600_ of 0.01 and grown in LB, and samples were harvested through the growth curve. The β-galactosidase assay was performed with ONPG (o-nitrophenyl-β-D-galactoside) as the substrate, and the results were calculated as Miller units normalized by OD_600_ (69).

### Fluorescence microscopy

T6SS firing events were detected by fluorescence microscopy using a VipA_sfGFP fusion separated by a 3XAla 3XGly linker, as previously described (17). Briefly, the *vipA* (without its stop codon) and *sfGFP* genes were amplified by PCR and cloned into the pJN105 vector to express a VipA_sfGFP fusion protein from an arabinose-inducible promoter. This construct was introduced by conjugation into the *C. violaceum* strains. The strains were grown under shaking in liquid LB at 37°C until midlog phase, when L-arabinose was added at a final concentration of 0.05% for 30 minutes prior to image acquisition. Cells were spotted in a thin pad of 1% agarose in PBS and imaged in a Carl Zeiss LSM 780 Microscope.

### Hcp cloning, expression, and purification

The *hcp* (CV_3977) coding region was amplified by PCR (Table S2) and cloned into the expression vector pET-15b. The construct was transformed into *E. coli* BL21(DE3). Recombinant His-tagged Hcp was induced by the addition of 1 mM isopropyl-D-thiogalactopyranoside (IPTG) to *E. coli* log-phase cultures. After 2 hours, the cells were harvested and lysed by sonication. His-Hcp was purified from the soluble fraction by NTA-resin chromatography (Qiagen). Purified protein was desalted with a PD-10 desalting column (GE Healthcare) in storage buffer (20 mM Tris-Cl, pH 7.4, 100 mM NaCl, 0.1 mM EDTA, and 5% (v/v) glycerol). Samples from the induction and purification steps were evaluated using 15% SDS-PAGE.

### Hcp detection in cellular and secreted fractions by western blot

*C. violaceum* strains were diluted to an OD_600_ of 0.01 and grown in liquid LB at 37°C until mid-log phase or the indicated time points throughout the growth curve. Two samples of each culture were harvested to obtain cellular and secreted fractions. For the cellular fractions, the samples were pelleted by centrifugation at 4,000 x g for 5 minutes. The secreted fractions were obtained by centrifugation at 4,000 x g for 15 minutes followed by filtration with 0.22 µm membranes to obtain cell-free supernatants. The pellets and supernatants were resuspended in SDS sample buffer to adjust the volumes according to the OD_600_ values of the cultures. The proteins were resolved by 15% SDS-PAGE and transferred to a nitrocellulose membrane. Western blotting was carried out with a Protein Detector™ LumiGLO^®^ Western Blotting (KPL) kit according to the manufacturer instructions. The polyclonal anti-Hcp antibody was used at a dilution of 1:10,000 for the cellular fractions and 1:2,000 for the supernatant fractions. Sample loading was assessed by membrane staining with 0.1% Ponceau S. The polyclonal anti-Hcp antibody was developed after immunization of six-week-old female BALB/c mice with three subcutaneous injections of the purified His-Hcp protein emulsified in Freund adjuvant (Sigma).

### VgrG3-HA coimmunoprecipitation (Co-IP) and mass spectrometry (MS)

To generate a VgrG3 protein fused at its C-terminus to a hemagglutinin tag (VgrG3-HA), the *vgrG3* gene containing its promoter and coding region but without its stop codon was amplified by PCR with specific primers. The long reverse primer contained the sequences for a duplicated HA tag and a stop codon (Table S2). The PCR-amplified fragment was digested with specific restriction enzymes and cloned into pMR20 (*vgrG*3-HA). This construct was introduced into the Δ*vgrG*3 mutant, generating the strain Δ*vgrG*3(*vgrG*3-HA). For the Co-IP assays, the WT(pMR20) as a control and the Δ*vgrG*3(*vgrG*3-HA) strains were diluted in LB for an OD_600_ of 0.01 and cultured until an OD_600_ of 0.8 was reached. The cultures (80 mL) were harvested by centrifugation, and the cells were resuspended in 7 mL of lysis buffer (150 mM NaCl, 10 mM Tris-Cl, pH 7.4, 1 mM EDTA, 1% Triton X-100, and 0.2 mM PMSF) prior to sonication. Lysed cells were treated with 1 ml 10% streptomycin for 20 minutes under agitation prior to centrifugation and filtration with a 0.45 µm filter. Proteins from the supernatant at 2 mg/mL were used for the Co-IP assay using Protein G magnetic beads (New England Biolabs) and a mouse monoclonal anti-HA antibody (Sigma). The anti-HA antibody was cross-linked to the Protein G magnetic beads using triethanolamine and ethanolamine. The protein samples were precleaned by incubation with Protein G magnetic beads. For Co-IP, cleaned samples (400 µg) were incubated with Protein G magnetic beads containing cross-linked anti-HA antibody for one hour at 4°C. The beads were washed three times with lysis buffer, and the proteins were eluted with elution buffer (0.1 M glycine, pH 2.5). For mass spectrometry, the eluted proteins were trypsinized, and the resulting peptides were analyzed by LC-MS/MS at the Life Sciences Core Facility (LaCTAD) from the State University of Campinas (UNICAMP).

## ACKNOWLEDGMENTS

This research was supported by grants from the São Paulo Research Foundation (FAPESP; grants 2020/00259-8 and 2021/06894-0) and Fundação de Apoio ao Ensino, Pesquisa e Assistência do Hospital das Clínicas da FMRP-USP (FAEPA). During the course of this work, JAA (grant 2018/03979-1) and MPM (grant 2018/12461-6) were supported by FAPESP fellowships, and FCL was granted a fellowship from Conselho Nacional de Desenvolvimento Científico e Tecnológico (CNPq). We thank the staff of the Life Sciences Core Facility (LaCTAD) from the State University of Campinas (UNICAMP) for the proteomics analysis and Roberta Rosales for technical support in microscopy.

## AUTHORS CONTRIBUTIONS

JAA, FCL, and JFSN planned the experiments; JFSN and JAA wrote the manuscript; JAA, FCL, and MPM performed the experimental work; JAA, FCL, and JFSN analyzed and interpreted the data; JFSN participated in study coordination and funding acquisition. All authors read and approved the final manuscript.

