## Supplementary material for "A quorum sensing-regulated type VI secretion system containing multiple nonredundant VgrG proteins is required for interbacterial competition in *Chromobacterium violaceum*"

### SUPPLEMENTAL MATERIAL

#### **Table S1.** Bacterial strains and plasmids

| Bacterial strain or plasmid | Description | Reference |
|---|---|---|
| ***Escherichia coli*** | | |
| DH5α | Strain for cloning purposes | (1) |
| S17-1 | Strain for plasmid mobilization | (2) |
| BL21(DE3) | Strain for protein expression | Novagen |
| ***Chromobacterium violaceum*** | | |
| ATCC 12472 | Wild type (Sequenced genome) | (3) |
| Δ*clpV* (CV_3965) | WT strain with CV_3965 gene deleted | This work |
| Δ*hcp* (CV_3977) | WT strain with CV_3977 gene deleted | This work |
| Δ*vipA* (CV_3979) | WT strain with CV_3979 gene deleted | This work |
| Δ*vgrG*1 (CV_3986) | WT strain with CV_3986 gene deleted | This work |
| Δ*vgrG*2 (CV_3975) | WT strain with CV_3975 gene deleted | This work |
| Δ*vgrG*3 (CV_1432) | WT strain with CV_1432 gene deleted | This work |
| Δ*vgrG*4 (CV_1233) | WT strain with CV_1233 gene deleted | This work |
| Δ*vgrG*5 (CV_0023) | WT strain with CV_0023 gene deleted | This work |
| Δ*vgrG*6 (CV_0016) | WT strain with CV_0016 gene deleted | This work |
| Δ*vgrG*1-2 | WT strain with CV_3986 and CV_3975 genes deleted | This work |
| Δ*vgrG*1-3 | WT strain with CV_3986, CV_3975 and CV_1432 genes deleted | This work |
| Δ*vgrG*1-4 | WT strain with CV_3986, CV_3975, CV_1432 and CV_1233 genes deleted | This work |
| Δ*vgrG*1-5 | WT strain with CV_3986, CV_3975, CV_1432, CV_1233 and CV_0023 genes deleted | This work |
| Δ*vgrG*1-6 | WT strain with CV_3986, CV_3975, CV_1432, CV_1233, CV_0023 and CV_0016 genes deleted | This work |
| Δ*cviR* | WT strain with CV_4090 gene deleted | (4) |
| Δ*cviI* | WT strain with CV_4091 gene deleted | (5) |
| WT(pSEVA) | WT strain containing the empty vector pSEVA | This work |
| Δ*vgrG*3(pSEVA) | Δ*vgrG*3 strain containing the empty vector pSEVA | This work |
| Δ*vgrG*3(*vgrG*3) | In trans complementation of *vgrG*3 with pSEVA vector in Δ*vgrG*3 | This work |
| WT(pMR20) | WT strain containing the empty vector pMR20 | This work |
| Δ*hcp*(*hcp*) | In trans complementation of *hcp* with pMR20 vector in Δ*hcp* | This work |
| Δ*vipA*(*vipA*) | In trans complementation of *vipA* with pMR20 vector in Δ*vipA* | This work |

| | | |
|---|---|---|
| Δ*vgrG*1-6(*vgrG*1) | In trans complementation of *vgrG*1 with pMR20 vector in the sextuple mutant (Δ*vgrG*1-6) | This work |
| Δ*vgrG*1-6(*vgrG*2) | In trans complementation of *vgrG*2 with pMR20 vector in the sextuple mutant (Δ*vgrG*1-6) | This work |
| Δ*vgrG*1-6(*vgrG*3) | In trans complementation of *vgrG*3 with pMR20 vector in the sextuple mutant (Δ*vgrG*1-6) | This work |
| Δ*vgrG*1-6(*vgrG*4) | In trans complementation of *vgrG*4 with pMR20 vector in the sextuple mutant (Δ*vgrG*1-6) | This work |
| Δ*vgrG*1-6(*vgrG*5) | In trans complementation of *vgrG*5 with pMR20 vector in the sextuple mutant (Δ*vgrG*1-6) | This work |
| Δ*vgrG*1-6(*vgrG*6) | In trans complementation of *vgrG*6 with pMR20 vector in the sextuple mutant (Δ*vgrG*1-6) | This work |
| WT(pCV_3981) | WT strain containing the promoter of CV_3981 with *lacZ* fusion | This work |
| Δ*cviR*(pCV_3981) | Δ*cviR* strain containing the promoter of CV_3981 with *lacZ* fusion | This work |
| WT(pCV_3982) | WT strain containing the promoter of CV_3982 with *lacZ* fusion | This work |
| Δ*cviR*(pCV_3982) | Δ*cviR* strain containing the promoter of CV_3982 with *lacZ* fusion | This work |
| WT(p*vgrG*1) | WT strain containing the promoter of *vgrG*1 with *lacZ* fusion | This work |
| WT(p*vgrG*2) | WT strain containing the promoter of *vgrG*2 with *lacZ* fusion | This work |
| WT(p*vgrG*3) | WT strain containing the promoter of *vgrG*3 with *lacZ* fusion | This work |
| WT(p*vgrG*4) | WT strain containing the promoter of *vgrG*4 with *lacZ* fusion | This work |
| WT(p*vgrG*5) | WT strain containing the promoter of *vgrG*5 with *lacZ* fusion | This work |
| WT(p*vgrG*6) | WT strain containing the promoter of *vgrG*6 with *lacZ* fusion | This work |
| Δ*cviR*(p*vgrG*3) | Δ*cviR* strain containing the promoter of *vgrG*3 with *lacZ* fusion | This work |
| Δ*cviR*(p*vgrG*4) | Δ*cviR* strain containing the promoter of *vgrG*4 with *lacZ* fusion | This work |
| WT(*vipA_sfGFP*) | WT with *vipA_sfGFP* cloned into pJN105 vector for L-arabinose induction | This work |
| Δ*clpV*(*vipA_sfGFP*) | Δ*clpV* with *vipA_sfGFP* cloned into pJN105 vector for L-arabinose induction | This work |
| Δ*hcp*(*vipA_sfGFP*) | Δ*hcp* with *vipA_sfGFP* cloned into pJN105 vector for L-arabinose induction | This work |
| Δ*vgrG*3(*vipA_sfGFP*) | Δ*vgrG*3 with *vipA_sfGFP* cloned into pJN105 vector for L-arabinose induction | This work |
| Δ*vgrG*1-3(*vipA_sfGFP*) | Δ*vgrG*1-3 with *vipA_sfGFP* cloned into pJN105 vector for L-arabinose induction | This work |

| | | |
|---|---|---|
| Δ*vgrG*1-6(*vipA_sfGFP*) | Δ*vgrG*1-6 with *vipA_sfGFP* cloned into pJN105 vector for L-arabinose induction | This work |
| Δ*cviR*(*vipA_sfGFP*) | Δ*cviR* with *vipA_sfGFP* under L-arabinose induction | This work |
| Δ*cviI*(*vipA_sfGFP*) | Δ*cviI* with *vipA_sfGFP* under L-arabinose induction | This work |
| Δ*vgrG*3(*vgrG*3-HA) | Δ*vgrG*3 in trans complemented with pMR20 expressing *vgrG*3 gene fused to hemagglutinin tag at C-terminal portion | This work |
| **Plasmids** | | |
| pNPTS138 | Suicide vector containing oriT, $Km^r$, *sacB* | D. Alley |
| pJN105 | Broad-host-range vector, araC-PBAD cassete; $Gm^r$ | (6) |
| pMR20 | Broad-host-range low-copy vector containing oriT, $Tet^R$ | (7) |
| pET-15b | His-tagged protein expression vector; $Amp^r$ | Novagen |
| pGEM-T easy | Cloning plasmid; $Amp^R$ | Promega |
| pRK*lacZ*290 | Vector containing promoterless *E. coli lacZ.* $Tet^R$ | (8) |
| pSEVA221 | Broad-host-range vector, $Km^R$, *oriRK2, oriT.* | (9) |

**Table S2.** Primers used in this work

| **Name** | **Sequence 5' - 3' *** | **Description** |
|---|---|---|
| **Construction of mutant strains** | | |
| CV_3965 del1<br>CV_3965 del2 | GATCATAAGCTTGATTGATGGGGCGTGGGAAG<br>GATCATGAATTCCGTCAACCGGTCTATCAGGG | 767 pb, *HindIII*/*EcoRI*. CV_3965 deletion |
| CV_3965 del3<br>CV_3965 del4 | GATCATGAATTCCGTATGGCCAGCGGCAAACC<br>GATCATGTCGACGCTTCAGTTCCTTGACCCGG | 652 pb, *EcoRI*/*SalI*. CV_3965 deletion |
| CV_3977 del1<br>CV_3977 del2 | GATCATAAGCTTGAAGTGGCGCTGAAGTGCCC<br>GATCATCTGCAGGCTTTCGCCAGGAATGCCATC | 633 pb, *HindIII/PstI*. CV_3977 deletion |
| CV_3977 del3<br>CV_3977 del4 | GATCATCTGCAGGGCAATGTCAGCGCCGGTTG<br>GATCATGGATCCGAGAGCTAAGGTCCATGGCG | 687 pb, *PstI/BamHI*. CV_3977 deletion |
| CV_3979 del1<br>CV_3979 del2 | GACTATGGGCCCATTCCCTGTCGCGCAGCCTG<br>GACTATGGATCCGCCGATCTCGACGTCATAGG | 607 pb, *ApaI/BamHI*. CV_3979 deletion |
| CV_3979 del3<br>CV_3979 del4 | GACTATGGATCCGACGAAGTGGTGCGCGATAC<br>GACTATGAATTCCCCACCAGCGCGGCGTAAG | 547 pb, *BamHI/EcoRI*. CV_3979 deletion |
| CV_3986 del1<br>CV_3986 del2 | TACCGGAAGCTTTGTACTTCGCGCTATTGGCC<br>TACCGGGGATCCGCTGGTCAGATCCATGATCG | 606 pb, *HindIII/BamHI*. CV_3986 deletion |
| CV_3986 del3<br>CV_3986 del4 | TACCGGGGATCCCTGAATGCGATCAAATCCGGC<br>TACCGGGAATTCTTGGCCAATGCATCGGGCTC | 663 pb, *BamHi/EcoRI*. CV_3986 deletion |
| CV_3975 del1<br>CV_3975 del2 | TACCGGAAGCTTGCCCCGGGCGGGTTTTTAAG<br>TACCGGGGATCCGTCCATGGCGCAATCTATCCG | 598 pb, *HindIII/BamHI*. CV_3975 deletion |
| CV_3975 del3<br>CV_3975 del4 | TACCGGGGATCCGTGGCCGCCAAATGAAGCCG<br>TACCGGGAATTCCCAACAGGTGGTTTTGCCGC | 660 pb, *BamHI/EcoRI*. CV_3975 deletion |
| CV_1432 del1<br>CV_1432 del2 | TACCGGAAGCTTCGGCCGCCAAACTACTAGTC<br>TACCGGGGATCCGTCAAGGTCCATCGATTGTCC | 626 pb, *HindIII/BamHI*. CV_1432 deletion |
| CV_1432 del3<br>CV_1432 del4 | TACCGGGGATCCTTCCCCTTCGTAGAGCAGCC<br>TACCGGGAATTCCGGCAGGAACGACGCCAATC | 603 pb, *BamHI/EcoRI*. CV_1432 deletion |
| CV_1233 del1<br>CV_1233 del2 | TACCGGAAGCTTCCAGGCTTCCAGATACTGCG<br>TACCGGGGATCCACTGGCTAGCAATGCGTTCAG | 611 pb, *HindIII/BamHI*. CV_1233 deletion |
| CV_1233 del3<br>CV_1233 del4 | TACCGGGGATCCCATTTCAACATGGTGCCGTCG<br>TACCGGGAATTCCCCTCCAGCGACTTCATTTC | 583 pb, *BamHI/EcoRI*. CV_1233 deletion |
| CV_0023 del1<br>CV_0023 del2 | TACCGGAAGCTTTGGCATGTTGGCAACAGGCG<br>TACCGGGGATCCCAGATCCATCGGCGTTCCAG | 593 pb, *HindIII/BamHI*. CV_0023 deletion |
| CV_0023 del3<br>CV_0023 del4 | TACCGGGGATCCCCGACGCTCCCCAAAGACAA<br>TACCGGGAATTCTGCATAGAACGGTGTGCCGG | 657 pb, *BamHI/EcoRI*. CV_0023 deletion |
| CV_0016 del1<br>CV_0016 del2 | TACCGGGGGCCCGAGATGCTAAGGCCGGTGAC<br>TACCGGGGATCCGAAGCTGGAGAGGAGGCTGT | 592 pb, *ApaI/BamHI*. CV_0016 deletion |
| CV_0016 del3<br>CV_0016 del4 | TACCGGGGATCCGCGAGTACCAACTTGCCCTTC<br>TACCGGGTCGACCTCCCCATGAAATGGATGGCG | 632 pb, *BamHI/SalI*. CV_0016 deletion |
| **Construction of complemented strains** | | |
| CV_3977 Comp Fw<br>CV_3977 Comp Rv | GATCATCTGCAGGAAGCCAAGGCCAAGTACCC<br>GATCATGAGCTCCGCAAGACAAGGCGACAACC | 852 pb, *PstI/SacI*. Product with *hcp* gene and its promoter region |
| CV_3979 Comp Fw<br>CV_3979 Comp Rv | GACTATGAATTCGGGGCGGCAATTGAACGATC<br>GACTATGAGCTCCGATGATGCTGTCGAGCAGG | 699 pb, *EcoRI/SacI*. Product with *vipA* gene and its promoter region |
| CV_3979 Over Fw<br>CV_3979_sfgfp Rv | GACTATGAATTCGAACATCTTGTAGAGAAGAGCC<br>GACTATCTGCAGGGGGGGGGGCGGCGGCGCTGCT<br>CGCCTCCTTCTTTCT | 526 pb, *PstI/EcoRI*. *vipA* full gene plus 3 Gly x 3 Ala linker |
| sfGFP_Fw<br>sfGFP_Rv | GACTATCTGCAGATGTCGAAAGGAGAAGAACTGT<br>GACTATGAGCTCTTTATACAGTTCATCCATTCCATG | 714 pb, *PstI/SacI*. Product with super folder GFP for cloning with *vipA* into pJN105 |
| CV_3977 Exp Fw<br>CV_3977 Exp Rv | GATCACATATGGCTTTTGATGCATTTCTGAAAATC<br>GATCAGGATCCTTAGGCGATTACTTTGTTCGCG | 498 pb, *NdelI/BamHI*. Product with *hcp* gene for heterologous expression |
| CV_3986 Comp Fw<br>CV_3986 Comp Rv | TACCGGGGTACCCTCGATCGAGCTGCATGAGC<br>TACCGGGGATCCCATCCCCCTTGTCCATTGCG | 2861 pb, *KpnI/BamHI*. CV_3986 complementation |
| CV_3975 Comp Fw<br>CV_3975 Comp Rv | TACCGGGGTACCCCGCCAATGTGCGCGATTAC<br>TACCGGGGATCCGGTGTCTTGCAACAGCATGC | 3019 pb, *KpnI/BamHI*. CV_3975 complementation |
| CV_1432 Comp Fw<br>CV_1432 Comp Rv | TACCGGGGTACCCGGGTCTCTCATGATCTTGCC<br>TACCGGGGATCCGCTATGGGTGCGGTCATGGC | 3554 pb, *KpnI/BamHI*. CV_1432 complementation |

| | | |
|---|---|---|
| CV_1432-HA_TagCompRv | TACCGGGGATCCTCAAGCGTAGTCTGGGACGTCGT ATGGGTAAGCGTAGTCTGGGACGTCGTATGGGTAG AACAGTTTGGGCAGGCTGGG | Primer used with CV_1432 Comp Fw. 3568 pb, *BamHI*. CV_1432 fused to HA epitope complementation |
| CV_1233 Comp Fw<br>CV_1233 Comp Rv | TACCGGGGTACCCCAGGCTTCCAGATACTGCG<br>TACCGGGAATTCGACGCGGTTTATGCTTGGCC | 3506 pb, *KpnI/EcoRI*. CV_1233 complementation |
| CV_0023 Comp Fw<br>CV_0023 Comp Rv | TACCGGGGTACCCGGTCCCAAGTCTTGCTGTG<br>TACCGGGGATCCCTCGGGAAGAGAACGGCATC | 2952 pb, *KpnI/BamHI*. CV_0023 complementation |
| CV_0016 Comp Fw<br>CV_0016 Comp Rv | TACCGGGGTACCGAGATGCTAAGGCCGGTGAC<br>TACCGGGGATCCCCGGCTCTCTACCAGTAATC | 3234 pb, *KpnI/BamHI*. CV_0016 complementation |
| **B-galactosidase assay** | | |
| pCV_3981Fw<br>pCV_3981Rv | GATCATGGATCCGCCAAGCACATTCTCGATTGG<br>GATCATAAGCTTGCGATTGTCGCCTGTTCCAC | 531 pb, *BamHI/HindIII*. Promoter region of CV_3981 for pRK*lacZ*290 cloning |
| pCV_3982Fw<br>pCV_3982Rv | GATCATAAGCTTGCCAAGCACATTCTCGATTGG<br>GATCATGGATCCGCGATTGTCGCCTGTTCCAC | 531 pb, *HindIII/BamHI*. Promoter region of CV_3982 for pRK*lacZ*290 cloning |

***** Underlined letters indicate the restriction enzyme recognition sites

**Table S3.** Bacteria strains used as prey in interbacterial competition assay

| **Bacteria** | **Strain** | **Abbreviation *** |
|---|---|---|
| *Burkholderia cepacia* | ATCC 17759 | Bc |
| *Escherichia coli* | ATCC 25922 | Ec |
| *Pseudomonas aeruginosa* | ATCC 27853 | Pa |
| *Staphylococcus aureus* | ATCC 29313 | Sa |
| *Salmonella typhimurium* | ATCC 14028 | St |
| *Stenotrophomonas maltophilia* | ATCC 13637 | Sma |
| *Enterobacter cloacae* | ATCC 13047 | Ecl |
| *Shigella flexneri* | ATCC 12022 | Sf |
| *Enterococcus faecium* | NCTC 13047 | Ef |
| *Klebsiella pneumoniae* | ATCC 13883 | Kp |
| *Klebsiella pneumoniae* | ATCC BAA-1705 | Kpn |
| *Enterococcus faecalis* | ATCC 4083 | Efa |
| *Shigella sonnei* | ATCC 25931 | Ss |
| *Pseudomonas aeruginosa* | PAO1 | Pao |

*Abbreviation referring to **Figure 2A**

**Table S4.** Proteins identified in coimmunoprecipitation assay of WT(pMR20) and Δ*vgrG*3(*vgrG*3-HA)

| **WT(pMR20)** | | | | | | |
|---|---|---|---|---|---|---|
| **Accession*** | **Description** | **mW (Da)** | **Peptides** | **Coverage (%)** | **Products** | **Amount (fmol)** |
| **Q7M7F1** | Elongation factor Tu | 43045 | 92 | 86,3636 | 974 | 32,4532 |
| Q7NQE7 | DNA-directed RNA polymerase subunit beta' | 155194 | 86 | 63,8252 | 640 | 2,0263 |
| **Q7NQF0** | Elongation factor G | 76958 | 60 | 79,7994 | 442 | 1,4946 |
| **Q7NQG4** | 50S ribosomal protein L5 | 20292 | 14 | 46,3687 | 70 | 0,3518 |
| **Q7NQG5** | 30S ribosomal protein S14 | 11589 | 7 | 38,6139 | 18 | 0,4214 |
| **Q7NQG8** | 50S ribosomal protein L18 | 12765 | 9 | 64,1026 | 52 | 0,5157 |
| **Q7NQG9** | 30S ribosomal protein S5 | 18207 | 14 | 47,6744 | 81 | 0,1538 |
| Q7NQM5 | Aspartate ammonia-lyase | 50287 | 24 | 76,8737 | 218 | 0,9885 |
| **Q7NQX1** | 60 kDa chaperonin 2 | 57382 | 40 | 69,7802 | 347 | 1,5874 |
| Q7NR97 | Nudix hydrolase domain-containing protein | 23223 | 10 | 66,8269 | 105 | 0,1679 |
| **Q7NUY8** | Trigger factor | 48546 | 34 | 81,3793 | 262 | 1,2455 |
| Q7NV09 | Uncharacterized protein | 6615 | 3 | 68,3333 | 38 | 0,0807 |
| Q7NV22 | Probable transmembrane protein | 13147 | 4 | 71,7742 | 53 | 0,0143 |
| Q7NVZ4 | 30S ribosomal protein S2 | 27112 | 18 | 69,1358 | 111 | 1,3853 |
| Q7NWY4 | Phosphonate metabolism protein PhnH | 20435 | 6 | 54,2105 | 66 | 0,0398 |
| Q7NX50 | Probable transcriptional regulator_ MerR family | 15218 | 9 | 58,4615 | 100 | 0,0483 |
| Q7NZ92 | Uncharacterized protein | 10198 | 4 | 43,8202 | 40 | 0,5778 |
| **Q7P095** | ATP synthase subunit beta | 50024 | 40 | 81,0753 | 434 | 2,0939 |
| **Q7P097** | ATP synthase subunit alpha | 54676 | 36 | 69,0661 | 327 | 2,4705 |
| **Q7P0N9** | Acetyltransferase component | 56484 | 19 | 49,0975 | 210 | 1,8117 |
| Q7P0P0 | Pyruvate dehydrogenase E1 component | 99230 | 67 | 81,2852 | 584 | 2,3749 |
| **Δ*vgrG*3(*vgrG*3-HA)** | | | | | | |
| **Accession** | **Description** | **mW (Da)** | **Peptides** | **Coverage (%)** | **Products** | **Amount (fmol)** |
| **Q7M7F1** | Elongation factor Tu | 43045 | 54 | 72,2222 | 714 | 45,6856 |
| **Q7NQF0** | Elongation factor G | 76958 | 39 | 55,5874 | 313 | 2,9557 |
| Q7NQF9 | 50S ribosomal protein L16 | 15401 | 4 | 30,4348 | 49 | 5,6009 |
| Q7NQG2 | 50S ribosomal protein L14 | 13409 | 15 | 51,6393 | 89 | 3,7829 |
| **Q7NQG4** | 50S ribosomal protein L5 | 20292 | 16 | 65,3631 | 108 | 0,739 |
| **Q7NQG5** | 30S ribosomal protein S14 | 11589 | 5 | 45,5446 | 56 | 4,4619 |
| **Q7NQG8** | 50S ribosomal protein L18 | 12765 | 7 | 52,9915 | 88 | 9,9647 |
| **Q7NQG9** | 30S ribosomal protein S5 | 18207 | 18 | 59,8837 | 149 | 3,5319 |
| Q7NQH0 | 50S ribosomal protein L30 | 6780 | 5 | 37,7049 | 33 | 1,1832 |
| **Q7NQX1** | 60 kDa chaperonin | 57382 | 18 | 46,1538 | 164 | 2,9832 |
| Q7NQZ9 | VgrG1 (CV_3986) | 93995 | 27 | 41,7431 | 314 | 0,2691 |
| Q7NR08 | Hcp (CV_3977) | 17881 | 8 | 53,3333 | 81 | 0,9134 |
| Q7NR10 | VgrG2 (CV_3975) | 94022 | 20 | 35,8945 | 255 | 1,3435 |
| Q7NRL5 | 30S ribosomal protein S21 | 8490 | 2 | 28,5714 | 19 | 1,6007 |

| | | | | | | |
|---|---|---|---|---|---|---|
| Q7NRT4 | 30S ribosomal protein S9 | 14356 | 4 | 36,1538 | 44 | 2,1704 |
| Q7NRV5 | 30S ribosomal protein S16 | 9536 | 7 | 59,0361 | 84 | 8,6455 |
| Q7NU57 | Uncharacterized protein (CV_2846) | 7517 | 2 | 45,4545 | 58 | 77,9891 |
| **Q7NUY8** | Trigger factor | 48546 | 15 | 42,7586 | 187 | 2,7161 |
| Q7NY43 | VgrG3 (CV_1432) | 107856 | 29 | 42,1782 | 363 | 2,6193 |
| **Q7P095** | ATP synthase subunit beta | 50024 | 27 | 51,1828 | 324 | 3,949 |
| **Q7P097** | ATP synthase subunit alpha | 54676 | 18 | 33,8521 | 217 | 4,4851 |
| **Q7P0N9** | Acetyltransferase component | 56484 | 12 | 32,3105 | 147 | 1,1308 |
| Q7P238 | VgrG5 (CV_0023) | 94125 | 17 | 29,9771 | 247 | 1,2663 |

*Accessions highlighted in bold indicate proteins identified in both assays and are probably contaminant proteins with affinity to the magnetic beads.

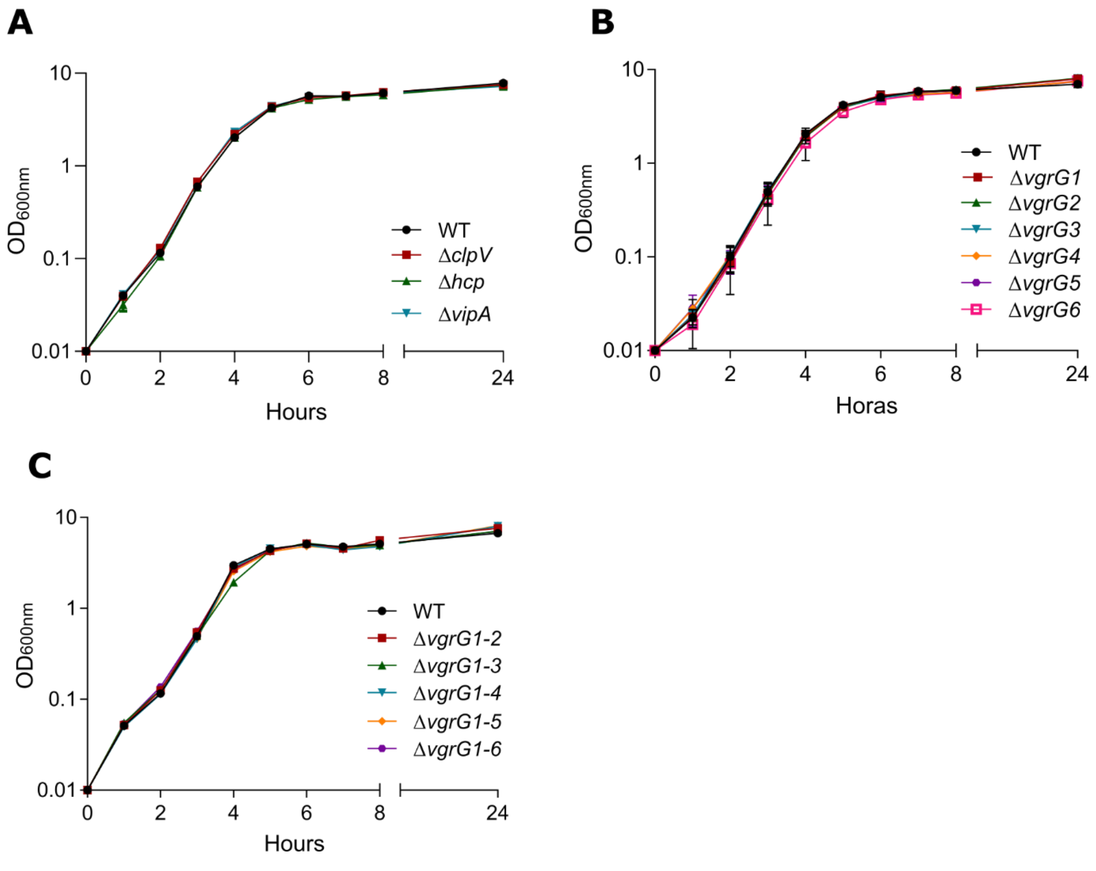


**Fig S1** Growth curves of the indicated strains grown in LB medium. (A) Mutants for T6SS core components. (B) Single mutant strains for each *vgrG*. (C) Sequential *vgrG* mutants. None of these strains showed any growth delay compared to the WT strain.

**A**

| | VgrG1 (CV_3986) | VgrG2 (CV_3975) | VgrG3 (CV_1432) | VgrG4 (CV_1233) | VgrG5 (CV_0023) | VgrG6 (CV_0016) |
|---|---|---|---|---|---|---|
| VgrG1 (CV_3986) | 100 | | | | | |
| VgrG2 (CV_3975) | 93.43 | 100 | | | | |
| VgrG3 (CV_1432) | 93.46 | 92.20 | 100 | | | |
| VgrG4 (CV_1233) | 76.46 | 76.23 | 71.06 | 100 | | |
| VgrG5 (CV_0023) | 84.36 | 84.27 | 83.55 | 82.50 | 100 | |
| VgrG6 (CV_0016) | 81.68 | 83.20 | 81.80 | 80.18 | 93 | 100 |

**B**

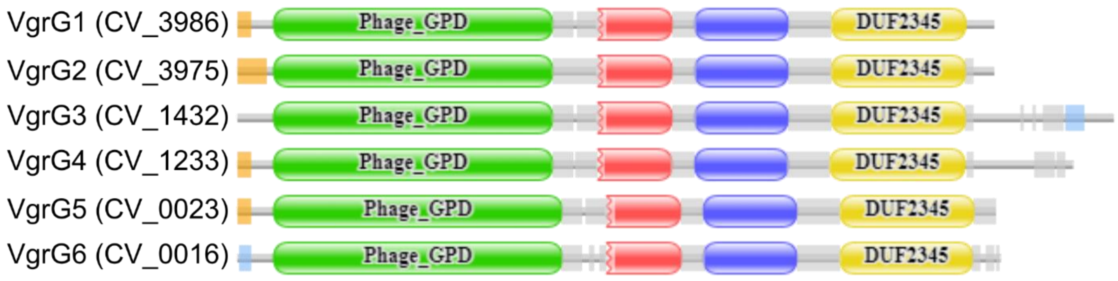


**Fig S2** The six VgrG proteins of *C. violaceum* show high sequence identity and similar domain organization. (A) Identity percentages shared among the six VgrG proteins of *C. violaceum*. The values were obtained with the Clustal Omega tool for multiple alignment. (B) Modular domains of VgrG proteins from analysis in the Pfam database. Green, red, and blue indicate the typical VgrG domains. All VgrGs have an additional C-terminal domain, DUF2345 (yellow). VgrG3 contains another additional region at C-terminus with low complexity (light blue). VgrG1, VgrG2, VgrG4, and VgrG5 have putative signal sequences at their N-termini (orange).
